# Rosace-AA: Enhancing Interpretation of Deep Mutational Scanning Data with Amino Acid Substitution and Position-Specific Insights

**DOI:** 10.1101/2025.01.09.632281

**Authors:** Jingyou Rao, Mingsen Wang, Matthew K Howard, Christian Macdonald, James S. Fraser, Willow Coyote-Maestas, Harold Pimentel

## Abstract

Proteins are dynamic systems whose function and behavior are sensitive to environmental conditions and often involve multiple cellular roles. Deep mutational scanning (DMS) experiments generate extensive datasets to capture the functional consequences of mutations. However, the sheer volume of data presents challenges in visualization and interpretation. Current approaches often rely on heatmaps, but these methods fail to capture the nuanced effects of amino acid (AA) substitutions, which are essential for understanding mutational impact. To address this, we extend the Rosace framework with `Rosace-AA`, a model that incorporates both position-specific information and AA substitution trends. Using substitution matrices like BLOSUM90, `Rosace-AA` offers a flexible and interpretable approach to summarize DMS data oil both protein-level and position-level. We demonstrate its utility across datasets, including OCT1 and MET kinase, showing that `Rosace-AA` highlights key positions where mutations deviate from expected substitution patterns and captures functionally relevant variation in protein behavior across multiple DMS screens. These results suggest that `Rosace-AA` enables more robust and interpretable analysis of complex DMS datasets.

## Background

In protein biology, a central question is whether a variant’s position within a protein or the type of amino acid substitution has a greater impact on function [1]. Research has shown that a variant’s position and substitution type can affect the mutation outcome.

Studies have demonstrated the importance of position, as certain amino acids play key roles in maintaining structural integrity, stability, or mediating interactions [2, 3, 4, 5]. For example, in G-protein-coupled receptors (GPCRs), some mutations can severely impair function due to their critical roles in ligand recognition and signal transduction [6, 7]. In contrast, mutations in surface-exposed loops are generally less consequential, as these regions are often peripheral to core functionality [6, 7] Other investigations highlight that the nature of the substitution itself is equally critical [8, 9]. Properties such as charge, hydrophobicity, size, and hydrogen-bonding potential could influence the functional impact of a mutation. Swapping amino acids with markedly different properties, such as changing a hydrophobic residue to a polar one, can disrupt function, especially if the original residue supports the protein’s core structure or active site [10, 11]. On the other hand, substitutions that maintain similar characteristics are often better tolerated.

Importantly, the interaction between the effects of position and substitution type also affects the outcome. A substitution’s impact often depends on its location within the protein structure: a change in a conserved and functionally critical region may lead to significant disruption, whereas the same substitution in a less crucial area might be more tolerable. This suggests that both factors matter, and that quantifying their respective impacts could provide a clearer understanding of mutation effects and relationships between position and substitution type.

Researchers have developed simple statistics to explore these questions. For example, using deep mutational scanning (DMS) experiments and calculating the average effect of mutations at each position or counting functionally significant variants offers insights into functional hotspots and sensitivity [12, 3, 13]. Others have examined amino acid substitution trends across positions, drawing on substitution matrices such as BLOSUM [10] and PAM [11]. However, a comprehensive, pattern-level analysis of the interactions between position and substitution type remains unexplored.

We extend the `Rosace` framework [14] to create a holistic framework which provides an intuitive interpretation of the interplay between position and substitution variant effects using DMS data. Our approach incorporates both position-specific information and amino acid substitution trends by decomposing variance into three components: (1) average effect at a position, (2) contribution of amino acid substitution, and (3) the remaining unexplained variation. This extended framework, termed `Rosace-AA`, provides three summary statistics for each position: the position-specific score, the position-specific scaling factor (susceptibility) for amino acid substitution effects, and the residual variance not explained by position or amino acid substitution. Those statistics quantify the effects of position, substitution, and their interaction, facilitating comparison within a protein and among experiments of the same protein under different conditions. For simplicity, we group amino acid substitutions based on BLOSUM90 in this paper, but the framework is flexible and can accommodate any user-defined substitution grouping.

Using `Rosace-AA`, we show that different protein domains exhibit distinct decomposition structures, as reflected by the relation of summary statistics across positions. Further, we validate the utility of these summary statistics by showing that predictions of protein secondary structure based on only the summary statistics achieve similar performance to the case when full DMS data are used. We demonstrate such utility by applying `Rosace-AA` to an OCTI DMS [12] to identify positions where variance deviates from global amino acid substitution expectations, emphasizing positions that disrupt local interactions and exhibit specific amino acid effects. Protein residues with high unexplained variance also suggest difficulty in prediction. Applying `Rosace-AA` to the MET dataset [3], a multi-screen analysis across 11 inhibitors and DMSO, it effectively summarizes multi-phenotype screens by identifying positions with distinct sensitivities or resistance profiles across various inhibitors. Unlike the OCT1 dataset, which focuses on differentiating positions based on alignment with global amino acid substitution expectations, the MET dataset application highlights `Rosace-AA`’s strength in capturing diverse phenotypic responses across multiple conditions, revealing functional hotspots with specific inhibitor interactions.

## Results

### Rosace-AA models both position and AA substitution effect

The `Rosace-AA` framework extends the original Rosace Bayesian hierarchical model [14] by modifying the prior for the variant-specific effect, denoted as *β*_*υ*_. In the original model, the prior mean of *β*_*υ*_ is represented by the position-specific effect *ϕ*_*p(υ)*_ in Figure 1A, where *p*(υ) is the mapping from a variant index to its corresponding position. This position-specific prior is shared across all variants at that location, which enforces regularization and variance shrinkage [14], in turn, increasing sensitivity and decreasing false discovery rate.

**Fig 1.**
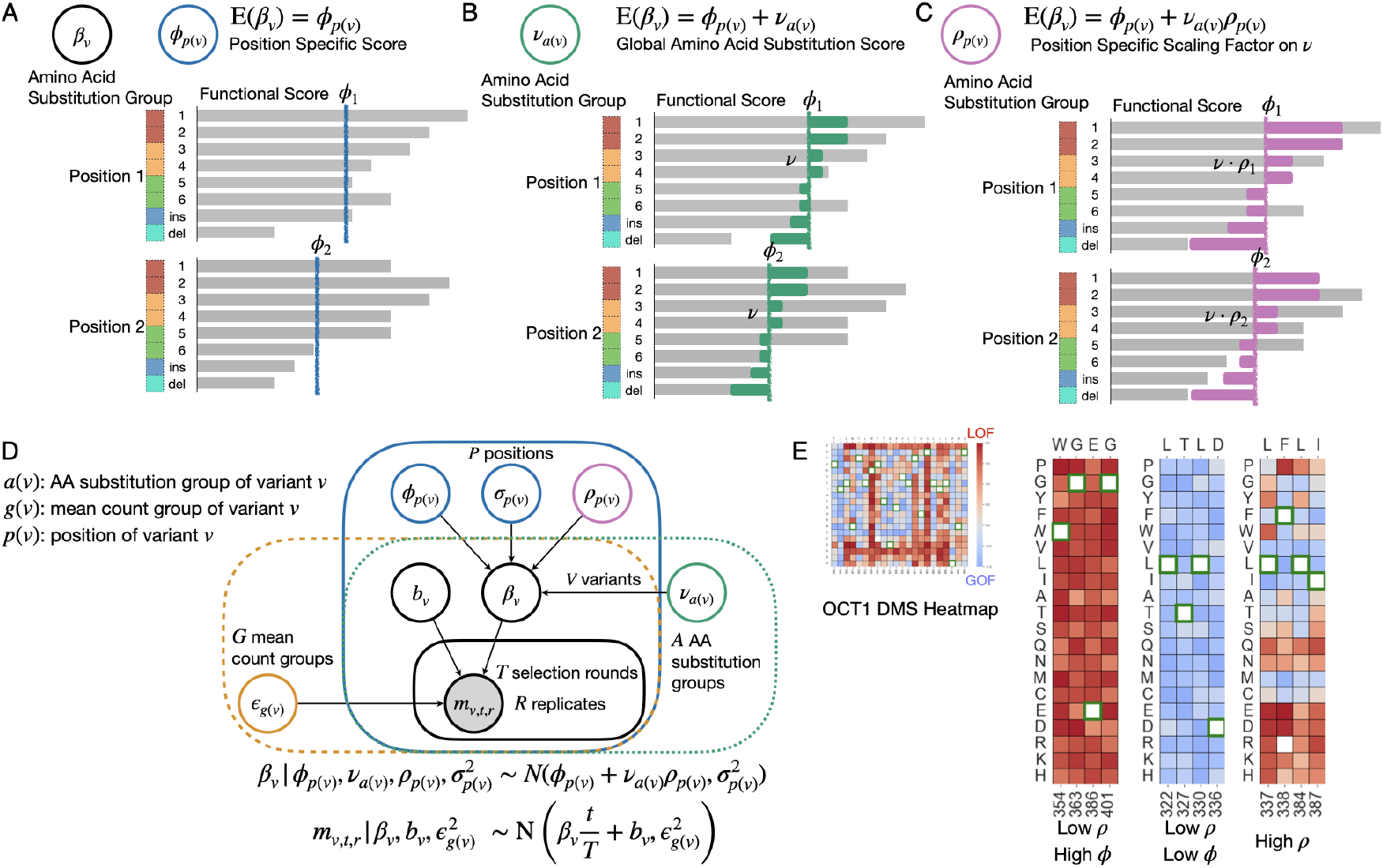
Overview of `Rosace-AA` model. A: Model 1 - original `Rosace` model. The prior or variant effect is the position-specific effect. B: Model 2 - `Rosace` with the addition of global AA effect prior. C: Model 3 - `Rosace` with the addition of position-scaled AA effect. D: Plate notation of Model 3. E: DMS functional score distribution for three types of positions, in the context of sensitivity to A A effects using OCTI dataset as an example.

In the extended model (Figure 1B), an additional term, *v*_*a*(*υ*)_is incorporated into the variant prior of *β*_*υ*_ to account for AA substitution effects, so that both the position and the substitution influence *β*_*υ*_. Here, *a*(υ) denotes the mapping from the variant index to its AA substitution group. As depicted in Figure 1B, the prior for the variant-specific score is now the sum of the position-specific score *ϕ*_*p(υ)*_ and the global AA substitution score *v*_*a(υ)*_ (represented by the end of the green bar). Variants with lower substitution scores, such as cysteine-to-glutamic acid mutations (green group in Figure 1B), are expected to exhibit a greater loss-of-function (LOF) effect compared to other variants at the same position. Conversely, variants with higher substitution scores, such as aspartic acid-to-glutamic acid mutations (red group in Figure 1B), are more likely to have a neutral or gain-of-function (GOF) effect. The substitution score thus indicates the sensitivity of a function score to AA substitution.

In this study, we employ a substitution matrix to group amino acids, restricting the AA substitution analysis to single missense mutations. Insertions and deletions across positions are grouped into a pseudo-position index, with no associated AA substitution effects. Similarly, synonymous mutations, which are pre-processed and normalized to an effect of approximately zero, are assigned a constant substitution effect of zero.

While there is a global overall impact of AA substitutions as observed in [10], we observe that the impact of AA substitution varies across positions. For instance, Figure 1E illustrates select positions from a deep mutational scanning experiment of the OCT1 cytotoxicity screen [12]. At some positions, the position-specific effect predominates, while others exhibit variable substitution effects. Based on this observation, we hypothesize that the AA substitution effects are heterogeneous across positions, prompting the introduction of an additional parameter, *p*_*p(υ)*_, referred to as the position-specific scaling factor, which ranges from 0 to 1 and modulates the influence of the AA substitution at a given position (Figure 1C). In this formulation, the prior for *β*_*υ*_ is now the sum of the positionspecific effect and the scaled AA substitution effect,*v*_*a(υ)*_ *ρ*_*p(υ)*_ · Figure 1C demonstrates that position 1 has a higher *ρ* value than position 2, allowing the prior mean of *β*_*υ*_ (the end of the purple bar) to better fit the data.

The complete plate diagram for this third model, is shown in Figure 1D. It is critical to note that all parameters of the prior for *β* _*υ*_ *—* namely *ϕ*_p*(υ)*_ *v*_*a(υ)*_ and *ρ*_*p(υ)*_ *—* are estimated jointly with *β*_*υ*_ and the rest of the model parameters. As a result, the estimation of *β*_*υ*_ is expected to have minimal bias from the experiment, and the regularization of the estimator through the prior mainly exerts on the uncertainty estimates of *β*_*υ*_,

### Variance decomposition of mutation effects

One key novelty of the `Rosace-AA` method lies in its ability to achieve variance decomposition through the prior specification of a target parameter (e.g. AA substitution effects) within a Bayesian hierarchical model. This method allows us to characterize how much of the variance in a mutation’s effect, within a protein or domain (depending on the scope of the DMS experiment), is attributable to different factors, such as position, AA substitution, and other unexplained sources (Figure 2B). Our approach extends the concept of variance decomposition, traditionally used in Analysis of Variance (ANOVA), by integrating it into the prior structure of a hierarchical model. Below, we explain the concept of variance decomposition in the context of ANOVA, highlight the key differences in our Bayesian formulation, and describe its application to our data.

**Fig 2.**
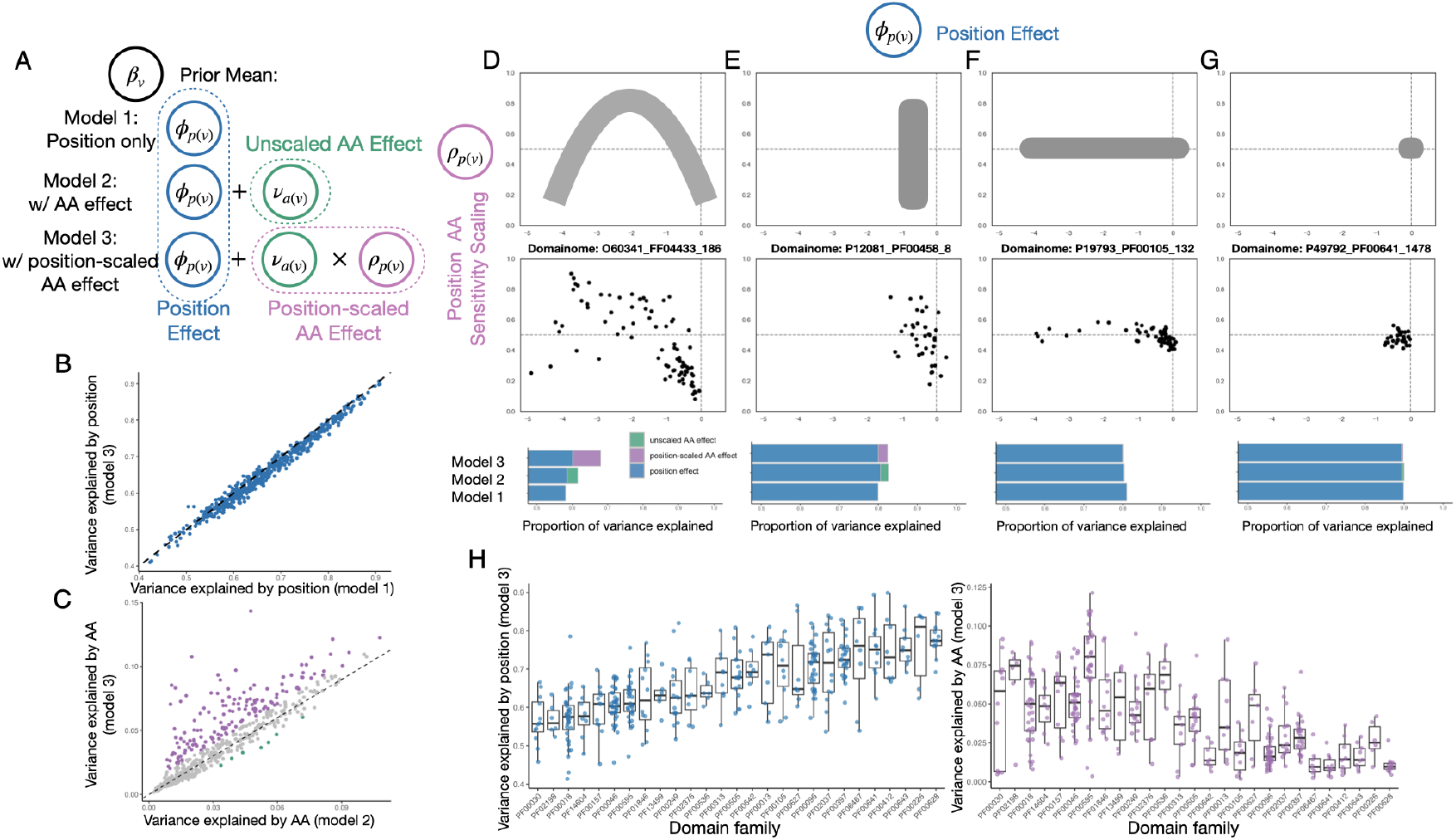
Global variance decomposition pattern and position-level effect trend applying `Rosace-AA` on Human Domainome 1.0 dataset. A: Model specifications revisited. B, C: Variance explained by position and AA components, compared across models. D-G: Top: the general shapes of the relation between position score and AA sensitivity scaling; Middle: an example of the relation described above; Bottom: Variance decomposition of each example domain. H: Variance explained by each factor by position, grouped by domain family.

Variance decomposition is commonly used in ANOVA. In the case of protein DMS data, we might consider the variance attributed to different positions within the protein sequence *p(υ)* (Model 1 in Figure 2A). In a one-way ANOVA, the goal is to compare the means of different positions and assess whether the observed differences are statistically significant. If a substantial portion of the total variance is attributed to between-group variance (i.e., between positions), we conclude that the position factor strongly influences the mutation effect.

However, our problem involves a more complex setting with two independent categorical factors: position *p*(*υ*) and AA substitution *a*(*υ*) (Models 2 and 3 in Figure 2A). Here, two-way ANOVA can be used to partition the variance into three components: the variance explained by position *ϕp*(*υ*) the variance explained by AA substitution *v*_*a(υ)*_ and the residual variance. This decomposition allows us to quantify the contributions of position and AA substitution to the overall variance in mutation effects.

Our model offers several advantages over ANOVA. First, while ANOVA aims to determine whether a factor significantly influences the target through a simple F-test, our model simultaneously performs multiple tasks. In our approach, *β*_*υ*_ the target parameter representing mutation effects, is treated as a random variable inferred from DMS count data, rather than being fixed and known *a priori*. This enables us to both estimate and perform hypothesis testing on *β*_*υ*_ Second, the factors *ϕ*_*p*(*υ*)_ and *v*_*a(υ)*_ are modeled as random effects, allowing the inference of their distributions. In contrast to a typical random-effects ANOVA, where the factor is treated as random but each group within the factor is assumed identically distributed, in our model each group within a factor has its unique posterior. Further, the uncertainty in estimating *β*_*υ*_ from the DMS count data is propagated into the variance estimates for each factor.

Finally, our model allows us to compute the percentage of variance explained by each factor and to characterize the mutation effect’s variance within a protein or domain. This decomposition provides insight into how much of the observed variance is due to position, AA substitution, or unexplained local factors, which can be summarized to characterize the protein. This rich variance decomposition serves as a powerful tool for understanding the role of different factors in protein function and evolution.

### Position effects dominate variance decomposition in Human Domainome data

We applied the three models described in Figure 1—Model 1 (Figure 1A), Model 2 (Figure 1B), and Model 3 (Figure 1C) — to over 500 human protein domains from the Human Domainome 1.0 dataset [15]. Note that this dataset directly measures domain stability, which is only a necessary condition for protein function. We computed the percentage of variance explained by position effects only (Model 1) and by both position and AA substitution effects (Models 2 and 3). Our analysis revealed that the variance explained by position is dominant in almost all proteins, with most proteins exhibiting a stability position effect ranging between 40% and 90%. Further, the variance explained by position remained consistent between Model 1 and Model 3 (Figure 2B), indicating the robustness of the model estimates despite the addition of an extra factor. This consistency also suggests that our method of decomposing the two factors is largely independent, even though position and amino acid substitution themselves are interdependent.

Our analysis identified that certain domain families consistently exhibit high variance explained by stability position effects. Notably, domain families such as PF00628 (PHD-finger), PF00226 (DnaJ domain), PF00643 (B-box zinc finger), and PF00412 (LIM domain) are among the top-performing groups, with position effects accounting for over 75% of the variance (Figure 2H). This suggests a strong influence of positional factors on these domains’ stability. In contrast, domain families (Beta/Gamma crystalline) and PF02198 (Ets-domain) rank among the lowest, with variance explained by position effects falling below 57% (Figure 2H). This lower value may indicate that positional effects play a less prominent role in these families or that other factors contribute more substantially to the variability.

In contrast, the variance explained by the global AA substitution effect, grouped by BLOSUM90, was relatively small, ranging from 0% to 10% in Model 2 (Figure 2C). However, the inclusion of the position-specific scaling factor in Model 3 increased the variance explained by the AA substitution effect for some proteins, with a few explaining approximately 10% of the total variance (annotated purple in Figure 2C). This demonstrates that while the overall contribution of the AA substitution effect is minor compared to the position effect, the position-specific interaction with AA substitutions can be significant for certain proteins. These results provide important insights into how both position and AA substitutions independently and jointly influence stability mutation effects within protein domains.

Protein domain families with higher variance explained by position tend to show lower variance explained by specific amino acids, which aligns with expectations. We focus on the PF00595 (PDZ domain) family for further analysis, as it exhibits a relatively narrow distribution in variance explained by position, yet a broad range in variance explained by amino acid substitutions. For instance, PDB domains on both DLG4 (P78352, 60) and MPDZ (075970, 372) show similar variance explained by position (60%). However, while DLG4 has no significant amino acid substitution effect, MPDZ shows over 12% variance explained by substitutions. This difference may indicate functional specialization or structural constraints within PDZ domains, where certain proteins might be more tolerant or dependent on specific amino acid changes, reflecting nuanced aspects of protein stability.

### Interaction between local position and AA substitution effect

In this section, we shift focus from the global protein-level summary statistics to an analysis of local, position-specific statistics. Conceptually, the protein or domain can be divided into four categories based on these plots: no effect, position-only, AA-substitution-only, and both. If the domain is mutation-insensitive, the points cluster near the origin (0,0) (Figure 2G). When position effects dominate but AA substitution has minimal impact, the points align horizontally at *ρ*_*p*(*υ*)_ = 0.5 (Figure 2F), suggesting uniform AA substitution effects across positions. Conversely, when position effects are neutral and AA substitution varies, the points form a vertical distribution (Figure 2E). In cases where both position and AA substitution effects vary, the scatter plot typically exhibits a “banana” shape (Figure 2D). Here, for strongly neutral or LOF positions, *ρ*_*p(υ)*_ is low, indicating that the position dominates regardless of the substitution. However, for mildly LOF positions, substitution plays a critical role, often depending on local structure and interactions with nearby residues, resulting in a curve that opens downward. While other shapes could theoretically emerge, these four categories capture the primary trends we observe.

These local statistics are directly produced by the `Rosace-AA` model. We extract two key position-level posterior distributions from the mutation effect prior. (1) the position-specific effect, *ϕ*_*p*(*υ*)_ ∈ (—∞, ∞), reflects the average mutation impact at a given position, with 0 indicating neutrality and more negative values signifying a stronger loss of function (LOF). (2) The position-specific AA substitution scaling factor, *ρ*_*p(υ)*_ ∈ [0,1], quantifies how much a position is affected by global AA substitution patterns. A *ρ*_*p(υ)*_ value of 0 indicates no global AA substitution effect, while a value of 1 reflects a significant influence. Further, since we employ a eta prior centered at 0.5 for *ρ*_*p(υ)*_ the model will default to 0.5 if the data lacks sufficient information (see Figure 2F-G). This behavior is observed either because (1) there is no global AA substitution effect, meaning *v*_*a(υ)*_ is 0, thus resulting in no overall effect, or (2) AA substitution effects are uniform across all positions, rendering scaling unnecessary.

Figures 2D-G demonstrates the different positional effects and AA substitution sensitivity patterns we have described. Each point represents a protein position, with the x-axis showing the position-specific effect and the y-axis representing the AA sensitivity scaling factor.

We further demonstrate the global protein-level summary statistics for the four categories, specifically, the proportion of variance explained by positional and AA substitution effects. In both the “no effect” and “position only” cases, positional effects dominate, explaining over 80% of the total variance. However, in cases where AA substitution effects are significant, we observe an increase in the overall proportion of variance explained, with AA effects contributing meaningfully to the model’s fit. Notably, in Figure 2D, the position-scaled AA effect (Model 3) provides a substantially better fit than the unscaled model (Model 2), highlighting the novelty and advantage of incorporating both positional scaled AA substitution effects in our approach. This result underscores the importance of modeling the interplay between position-specific and AA substitution effects for a more accurate representation of protein mutational landscapes.

### Position-specific statistics accurately predict secondary structure

To demonstrate the utility of the proposed position-specific summary statistics in capturing overall protein mutational landscapes, we applied them to a classical secondary structure prediction task using single missense deep mutational scanning (DMS) data. Suppose the performance of using only the summary statistics (*ϕ*_*p*(*υ*)_ and *ρ*_*p(υ)*_ 2 per position) is comparable to using the full DMS functional scores (*β*_*υ*_, 20 scores per position) (Figure 2B). In that case, this strongly supports that these position-specific statistics are effective and informative.

We selected secondary structure prediction for several reasons. First, it is a well-established problem so that it serves as a reliable validation task. Second, secondary structure prediction is critical for understanding broader protein structure and property prediction on a larger scale[16]. Finally, classical approaches to protein structure prediction, such as hidden Markov models and position-specific scoring matrices using multiple sequence alignment data, are position-informative. We therefore hypothesize that DMS data, which also contain rich positional information, should contribute valuable insights into secondary structure prediction.

To prevent data leakage, we use entirely distinct datasets for training and testing, incorporating an element of transfer learning. The training data consists of over 500 human protein domains (around 50 amino acids) from the Human Domainome 1.0 dataset [15], while the testing dataset is based on complete proteins or large domains (longer than 250 amino acids). While domains may appear in various proteins with different functions, they remain contextually distinct. This separation between domains and proteins reduces the potential for data leakage, creating a reliable validation process.

For the model architecture, we employ an ensemble of gradient boosting classifiers with different positional windows, combined with an ensemble layer to compute the weighted likelihood for secondary structure classification (Figure 3A; detailed in the Methods section). For simplicity, we only have three categories: *α*-helix, *β*-strand, and other. For benchmarking, we also applied a sequence-based second structure predictor `JPred` 4[17]. Note that its predictions are hard predictions instead of probabilities.

**Fig 3.**
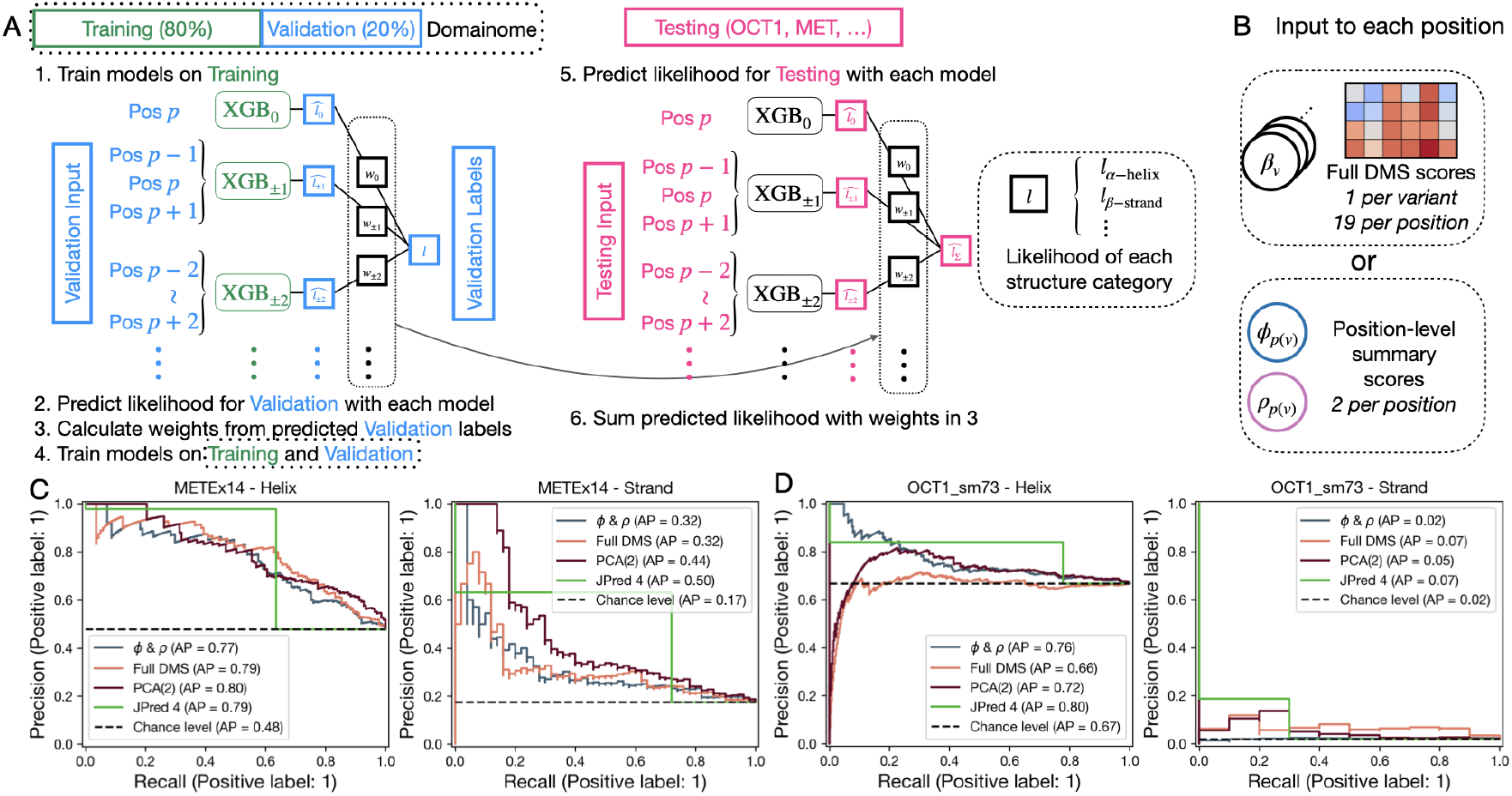
Position-specific statistics demonstrate utility in secondary structure prediction task. A: Steps of secondary structure prediction. B: Example inputs to the models. C, D: Recall-precision curves of predictions with different inputs and methods, applied to MET and OCTI.

We evaluated the model on the OCT1 cytotoxicity screen [12] and the MET kinase domain screen [18], both of which measure protein function, unlike the training and testing datasets which measure stability. The precision-recall curves indicate that using the full DMS scores, the first two principal components of full DMS scores, or the position-level summary statistics yield comparable results (Figure 2C and 2D), supporting that our summary statistics are robust. Analyzing the ensemble layer’s position window weights, we observed that the weights on selected sliding windows are distributed evenly, whether we use the summary statistics, full DMS scores, or their principal components.

### Unexplained mutation effects beyond position and global AA substitution require closer examination

The `Rosace-AA` model captures crucial information with the Bayesian hierarchical prior of functional score *β* _*υ*_ which is modeled as a normal distribution with both prior mean and prior variance (Figure 4A). Previously, we focused on the composition of the prior mean without leveraging the prior variance estimate 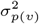 Here, we explore the residual variance — specifically, the portion of the score that is not explained by position or global amino acid (AA) substitution — since this residual component is likely to be less predictable and may reveal important insights.

**Fig 4.**
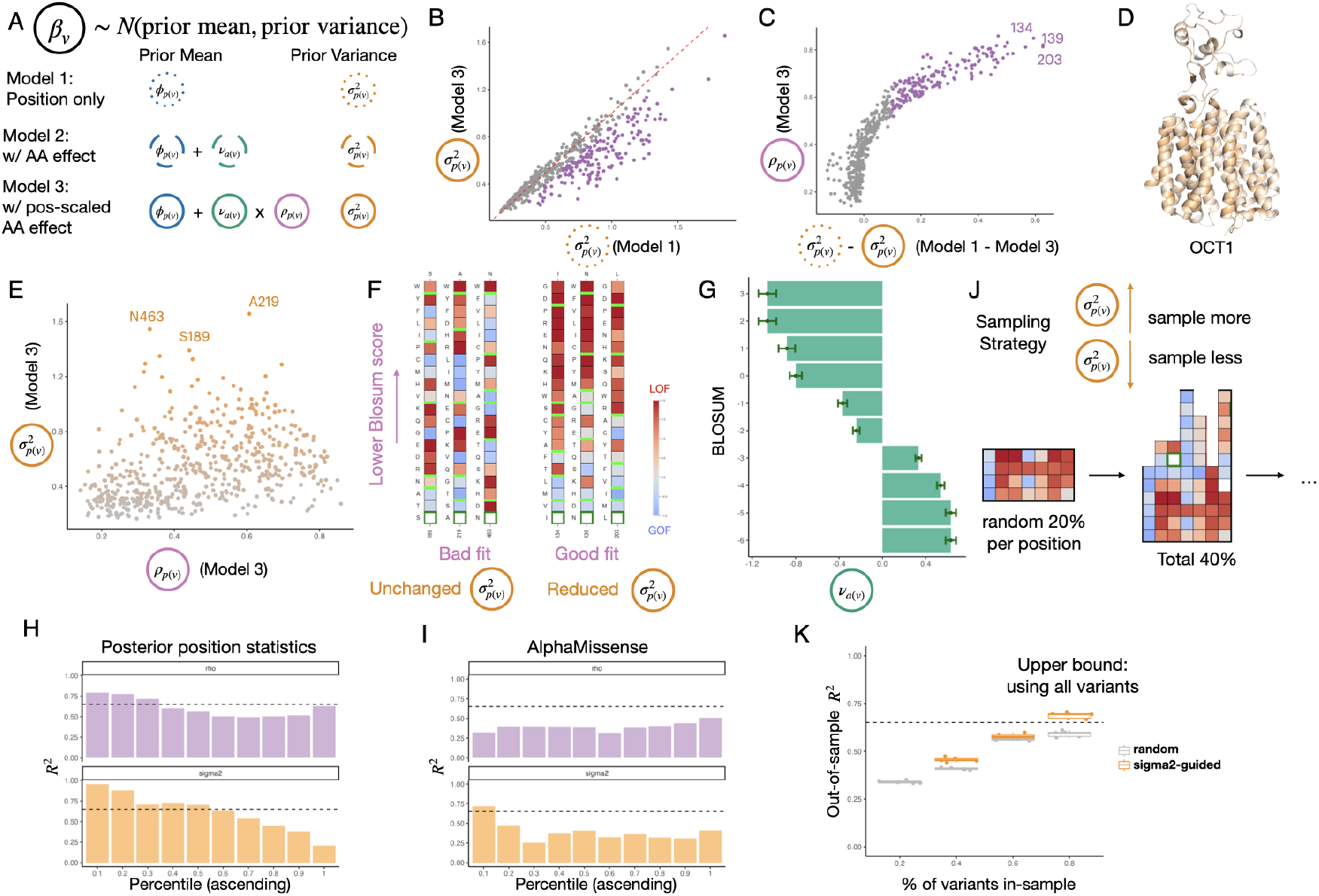
Applying `Rosace-AA` on OCTI dataset. A: Model specifications revisited. B: Unexplained variance in Models 1 and 3. C: Difference in unexplained variance is non-linearly attributable to A A substitution effects. D: Structure of OCTI, colored according to unexplained variance. E: Sensitivity to AA substitution and unexplained variance by position - the same color is used on D. F: Functional score heatmap of positions with good fit or bad fit under Model 3. G: Estimated AA substitution effect by BLOSUM group. H, I: Explained variance (R-squared) of each model, by *ρ* and *σ*^2^ quantiles. J: Variant sampling strategy. K: Difference in out-of-sample R-squared using random sampling or *σ*^2^-guided sampling.

The position-specific variance, 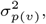 quantifies the unexplained variance in the variance decomposition outlined in Section 2.2 of the Results. With the inclusion of variance explained by AA substitution in Model 3 (Figure 4A), one would expect the unexplained variance to remain the same or decrease when a significant AA substitution effect is observed, indicated by a higher *ρ*_*p(υ)*_value (Figures 4B and 4C). This relationship is exemplified by the OCT1 cytotoxicity data [12].

To pinpoint the regions exhibiting the greatest unexplained variance in OCT1, we mapped 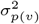 onto the OCTI structure (Figure 4D) and plotted it against the position-specific AA substitution scaling factor, *ρ*_*p(υ)*_. We expected an increase in variance explained by the global AA substitution matrix would lead to a decrease in 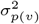 However, our observations contradicted this expectation, revealing a seemingly random pattern instead. Notably, the positions with the highest 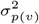 values (189S, 219A, and 463N) showed a broad range of *ρ*_*p(υ)*_ values. This unexpected finding prompts a closer investigation into the functional scores associated with these mutations.

To delve deeper into our analysis, we compared two distinct groups of positions: one exhibiting high 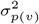 values, termed the “poor fit” group for global AA substitution, and another characterized by high *ρ*_*p(υ)*_ and low 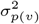 values, referred to as the “good fit” group. We hypothesized that positions with lower BLOSUM scores would exhibit more pronounced LOF effects, indicated by higher *v*_*a(υ)*_ values, while those with higher BLOSUM scores would show lesser LOF effects (Figure 4G). Our findings supported this hypothesis within the “good fit” group, where substitutions with lower BLOSUM scores indeed demonstrated more significant LOF effects, as expected (Figure 4F). Conversely, the “poor fit” group displayed erratic LOF effects that did not align with the anticipated BLOSUM trend. This discrepancy suggests that local factors, such as protein-protein interactions or conformational changes, may be influencing the functional impacts at these positions, rather than the overarching evolutionary or biophysical patterns associated with amino acid substitution. These positions are particularly compelling, as they may involve intricate local interactions within the protein or interactions with external factors, such as drugs, as seen in the OCT1 cytotoxicity screen.

### High positional unexplained variance suggests challenges in variant effect prediction

We hypothesized that positions with the largest unexplained variance, accounting for both position-specific and global AA substitution effects, would be more difficult to predict using variant effect predictors like AlphaMissense [19]. This also motivates experiment designs to sample more on such positions with high unexplained variance, to compensate for the diminished efficacy of statistical methods. Conversely, suppose an experiment is constrained on the number of variants to generate and test. In that case, researchers can devote more samples to positions with larger unexplained variance and use predictors to predict other variants’ effects more reliably.

To test the hypothesis that high unexplained variance on a position indicates difficulty in variant effect prediction, we approached the prediction task as a linear regression within a Bayesian hierarchical model, using the prior mean (Model 3, Figure 4A) derived from all deep mutational scanning (DMS) scores to predict variant effects. This setup represents the upper bound of prediction performance for a linear regression-based method, as it leverages all available data. We evaluated prediction accuracy using the R-squared metric, which reflects the proportion of variance explained by position and global AA substitution (Figure 4H). To identify which features are associated with greater prediction difficulty, we grouped variants into deciles based on position-specific AA substitution scaling factor *ρ*_*p(υ)*_ and variance 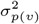.

As anticipated, when grouping variants by *ρ*_*p(υ)*_ the R-squared values remained relatively uniform across percentile groups, indicating that the model effectively captures AA substitution information, as specified by the substitution matrix. However, when grouping by 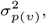 we observed that positions with higher variance were significantly harder to predict compared to those with lower variance, particularly for the highest 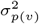 decile. This suggests that higher 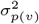 may indicate difficulty in variant effect prediction.

We then analyzed predictions from AlphaMissense. Interestingly, positions with higher *ρ*_*p(υ)*_ values showed a slight improvement in prediction accuracy, suggesting that evolutionary trends, captured by substitution matrices like BLOSUM, may enhance predictive performance (Figure 4I). Regarding 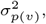 the lowest 10% of positions exhibited high R-squared values, approaching 75%, while the highest variance positions saw this metric drop to around 30%. These findings support our initial hypothesis: positions with higher unexplained variance from DMS are indeed more difficult to predict.

Finally, we explored the potential for improving variant effect predictions in scenarios where complete deep mutational scanning (DMS) experiments are infeasible — such as for large proteins or other experimental constraints — by employing a more efficient sampling strategy guided by estimates of positional variance 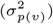. The guiding principle is straightforward: when positional variance is high, more variants are sampled from that position, and when variance is low, fewer variants are sampled (Figure 4J). To test this, we devised a toy example where, in the first round, we randomly sampled 20% of variants at each position. In the second round, we sampled another 20 either completely at random or guided by 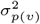. We then used the parameters learned from these sampled datasets with the `Rosace-AA` model to predict, the functional scores of the remaining, unsampled variants, using R-squared as the evaluation metric. As shown in Figure 4K, the 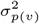-guided sampling strategy resulted in a notable improvement in prediction accuracy compared to random sampling, demonstrating the effectiveness of variance-guided sampling in enhancing prediction performance.

### Example: Rosace-AA for multi-phenotype DMS analysis

To demonstrate the utility of `Rosace-AA` in multi-phenotype analysis, we applied the model to data from a DMS experiment on the MET kinase domain under various ATP-competitive inhibitor conditions [3].

MET is a Receptor Tyrosine Kinase (RTK) involved in cellular growth and survival, and its dysregulation — through mutations or amplification — has been implicated in cancers such as lung and gastric cancers. Targeting MET with inhibitors blocks aberrant signaling pathways contributing to cancer progression. MET utilizes ATP for signal transduction, and many inhibitors compete with ATP binding to inhibit its kinase activity. However, mutations in MET can lead to resistance against specific inhibitors, necessitating the evaluation of multiple inhibitors to determine the most effective treatment [3].

In this DMS dataset, experiments were conducted under a control condition and with 11 different ATP-competitive inhibitors. For each condition, we separately computed the variant-level fitness scores (*β*;_*υ*_) and two position-level summary scores, *ϕ*_*p*(*υ*)_ and *ρ*_*p(υ)*_, using `Rosace-AA` (Figure 5A).

**Fig 5.**
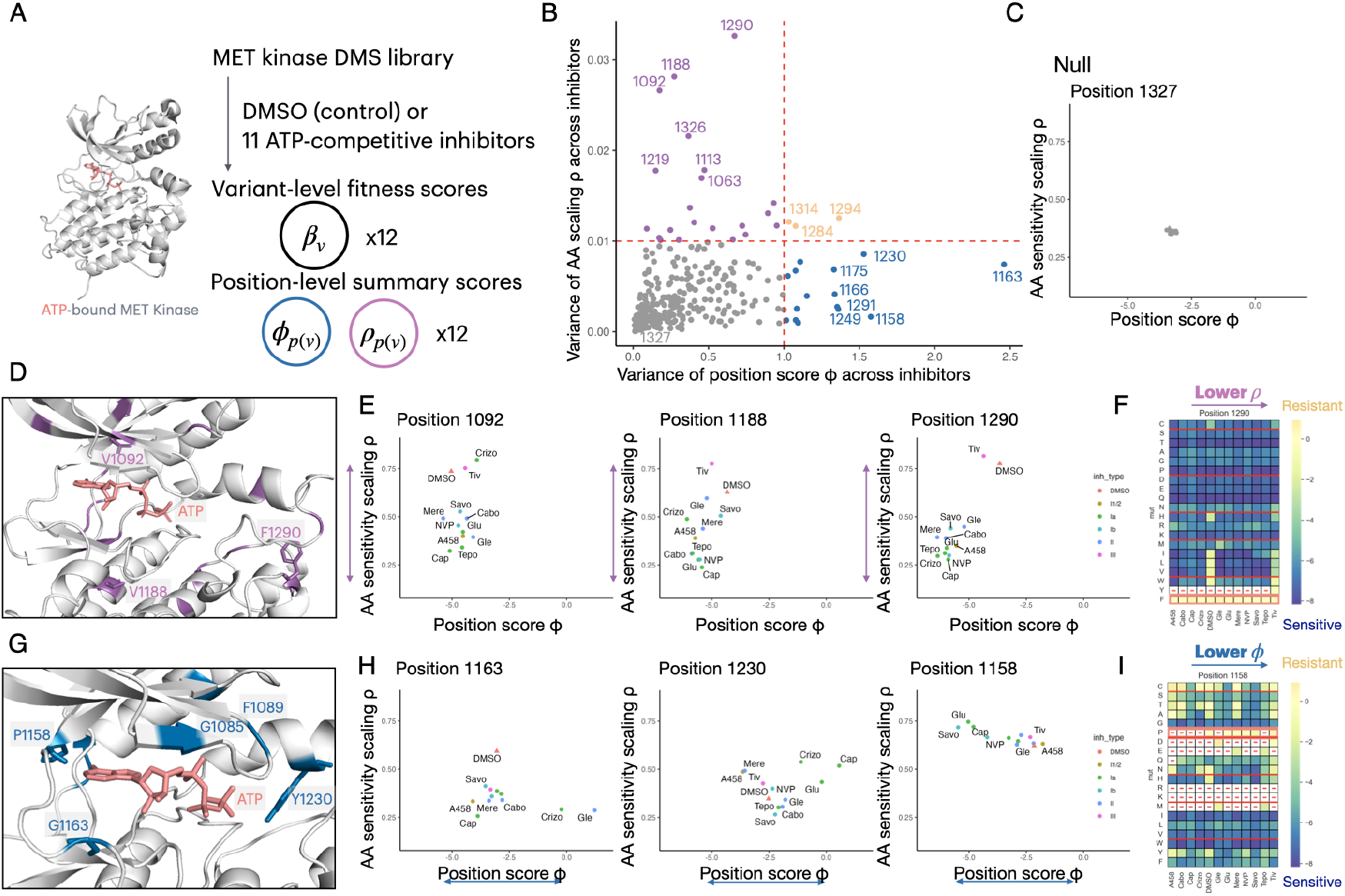
Applying `Rosace-AA` on MET kinase domain DMS dataset. A: Experiment description. B: Cross-inhibitor variance of *ϕ* and *ρ* by position, colored according to cross-inhibitor consistency of each factor. C: Example of both *ϕ* and *ρ* being consistent on a position across inhibitors. D, E: Examples of consistent *ϕ* but varying *ρ* positions. F: Functional score heatmap of a position with consistent *ϕ* but varying *ρ*. G, H: Examples of consistent *ρ* but varying *ϕ* positions. I: Functional score heatmap of a position with consistent *ρ* but varying *ϕ*.

To identify positions with inhibitor-specific effects, we computed the variance of these position-level scores and visualized them in a scatter plot. Positions with minimal variability across inhibitors represent the null condition, such as position R1327, which showed consistent behavior across all inhibitors (Figure 5C). Positions of greater interest exhibit significant variance in either *ϕ*_*p*(*υ*)_ (blue in Figure 5B) or *ρ*_*p(υ)*_ (purple in Figure 5B), indicating inhibitor-specific effects. We mapped these key positions onto the MET structure (Figures 5D and 5G).

The top three positions with high variance in *ϕ*_*p*(*υ*)_ were located within the ATP-binding pocket of MET (Figure 5G), suggesting that mutations at these sites produce inhibitor-specific effects at the inhibitor-binding pocket without broadly affecting overall protein function. Each of these three positions displayed distinct resistance profiles to the different inhibitors (Figure 5H). For example, MET inhibitors can be grouped by their binding mode to the ATP pocket and conformational preference [3]. Mutations to Position P1158 are more sensitive to Types I1/2, II, and III inhibitors and more resistant to Types Ia and Ib inhibitors (this is also shown in Figure 5I), implying that the mutations are likely blocking the binding of the latter inhibitors and facilitating that of the former ones. Position Y1230 displays a roughly opposite profile. Position G1163 is the most interesting case among the three as sensitivity to inhibitors is apparently not related to inhibitor type.

In contrast, the top three positions with high variance in *ρ*_*p(υ)*_ were located farther from the ATP-binding pocket, suggesting a more distal allosteric effect (Figure 5D). Interestingly, these positions shared a common substitution profile: DMSO and Tivantinib (Tiv) inhibitors displayed high general amino acid substitution effects, while Capmatinib (Cap) showed the lowest (Figure 5E). Further inspection of other positions with high *ρ* shows that Tiv generally has similar behavior to that of DMSO, in contrast to other inhibitors, probably due to its MET independence and a different inhibition mode. A detailed examination of the heatmap for position V1290 revealed that substitutions to Ile, Leu, Val, and Trp were driving these changes of *ρ*_*p(υ)*_ (Figure 5F).

In summary, this MET analysis demonstrates how Rosace-AA’s position-level summary statistics can reveal nuanced insights into the functional impact of mutations across different phenotypic conditions. By capturing variance in mutation effects both locally (within critical functional sites) and distally (at allosteric positions), Rosace-AA enables the identification of mutation-specific and condition-specific functional shifts. This approach not only aids in understanding the mechanistic underpinnings of inhibitor resistance but also provides a broader framework for multi-phenotype analyses, allowing researchers to explore complex genotype-phenotype relationships in various experimental contexts.

## Discussion

The `Rosace-AA` framework provides a new powerful approach to dissecting the functional effects of mutations across diverse conditions by incorporating both position-specific information and amino acid substitution trends. Its ability to classify positions based on mutation effects and susceptibility to amino acid changes has broad applications in fields such as precision medicine, protein engineering, and drug discovery. For example, the insights gained from the MET DMS dataset highlight its potential for identifying mutation-driven resistance mechanisms in cancer therapeutics. By extending this framework to other proteins or drug-target interactions, researchers can uncover novel functional insights that inform personalized treatment strategies or guide the design of more effective inhibitors.

A current major challenge in DMS is to gain biological insight from the screen. It usually involves ad hoc quantitative analyses of sequence patterns and protein structure. Incorporating mutational information allows for identifying positions that statistically diverge from the background. Divergent variants and their residues are likely to have distinct roles in protein function such as binding interfaces, catalytic sites, and other crucial positions, and thus merit further inspection.

Despite its strengths, the `Rosace-AA` framework has limitations. First, the decomposition of variance relies on predefined substitution groupings, such as the BLOSUM90 matrix, which may not fully capture all biological contexts. The framework’s effectiveness in analyzing other protein families with distinct evolutionary or functional constraints remains to be thoroughly tested.

Future research could extend `Rosace-AA` by incorporating more sophisticated models of amino acid substitution, allowing users to input custom substitution matrices. This flexibility would enhance the framework’s applicability across diverse protein domains and evolutionary contexts, providing more biologically relevant insights. Another important direction is to model epistatic interactions with multi-mutation DMS data, enabling `Rosace-AA` to capture non-linear mutation effects that go beyond position-specific and substitution effects. This extension would provide a more comprehensive view of mutational impacts, especially in cases where substitution patterns are more complex. Additionally, the positional features extracted by `Rosace-AA` could guide more targeted subsampling of mutations, facilitating the mapping of mutational landscapes and helping prioritize regions of interest for further experimental or computational study.

## Methods

### Rosace-AA: Raw DMS Read to Sequencing Count

Following the notation of Rosace, in a growth-based DMS screen, each raw sequencing count is denoted by *c*_*υ*_,_*t,r*_, standing for variant, selection round, and replicate indices respectively.

*p*(·) encodes the positional information of a variant. In most cases, it maps a variant to its amino acid position. There are two notable exceptions, however: (1) If information is given whether each variant is synonymous or not, all synonymous variants are grouped as a virtual “control” position. (2) If a position has too few variants (due to missing data, for example), all variants of this position will be merged into the next position. The process is repeated if the combined position also lacks sufficient variants.

*u*(·) is a similar mapping that extracts the mutant corresponding to a variant, it can be: (1) substitution (e.g. RHK → RDK), (2) insertion (e.g. RHK → RHDK), or (3) deletion (e.g. RHK → RK). In this paper, the terms “mutant” and “AA substitution” are used interchangeably.

Every possible mutant is further assigned a mutant group label, with *A* possible mutants in total, *a*(·) thus maps variants to such “AA substitution groups”. Since each AA substitution group contains multiple mutants, *a*(-) is by construction a coarser mapping than *u*(·).

### Rosace-AA: Preprocessing of Sequencing Counts

`Rosace-AA` preprocesses raw sequencing counts using the same sequence of steps as used in `Rosace`:

1. Variant filtering: For each replicate, if a variant has more than 50% missing count data, it will be dropped.
2. Missing data imputation: If a variant has fewer than 50% of missing counts, they are imputed using the K-nearest neighbor averaging (*K* = 10) or filled with 0.
3. Scale transformation: The imputed counts are offset by 1/2 (pseudo-count) and log-transformed.
4. Count normalization: The transformed counts are normalized by either the wild-type sequences or the sum of sequencing counts for synonymous mutations. This approach was proposed by *Enrich2* [20] to approximately center the wildtype functional score to 0. Additionally, the pre-selection (*t* = 0) count is subtracted to simplify the specification of the intercept’s prior.

The resultant aligned count is identified by *m*_*υ t, r*_ with *υ, t*, and *r* defined above. Scale transformation and count normalization are as follows:

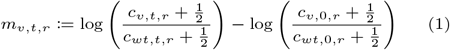

### Rosace-AA: Bayesian Hierarchical Model

Following the convention of Rosace, Rosace-AA assumes linear growth of the aligned count *m*, with regard to time *t* (selection round), expressed in mathematical terms:

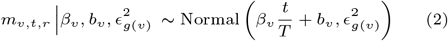

where the growth rate *β*_υ_ is treated as the functional score of the variant and *b*_*υ*_ as the intercept. *ϵ*_*g(υ)*_ is the scale of the error, grouped by the mean group of the variant *g*(υ).

Additionally, the mean variant-level growth rate *β*_υ_ is modeled as follows (terms explained below):

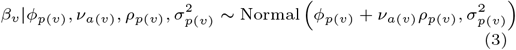

### Position-level growth rate and error

`Rosace`, the predecessor of `Rosace-AA`, models the mean variant-level growth rate *β*_υ_ with a mean effect *ϕp*(*υ*) and an error of magnitude 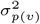 both of which are position-dependent:

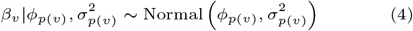

It follows the assumption that non-synonymous variants at the same position have similar effects and a similar scale of errors. Note that synonymous mutations are assigned a virtual control position and positions with an insufficient number of variants are merged.

The position-level mean effect *ϕ* and variance *σ*^2^ are, consistent with `Rosace`, given weakly-informative priors: *ϕp*(*υ*) ∼ Normal(0,1) and 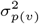 ∼ InvGamma(1, 1)

### A A-substitution-level growth rate and activation

`Rosace-AA` improves upon Rosace by incorporating AA-substitution-level information. It has two additional components to the modeling of *β*_υ_ as in Equation 3: (1) AA-substitution-group-level functional effect *v*_*a(υ)*._ It can be loosely interpreted as the functional “potential” inherent to an AA substitution. This term is not directly additive with *ϕp*(*υ*), but regulated by: (2) Position-level activation score *ρ*_*p(υ)*_. The AA substitution has the largest impact at positions with the highest activation scores (the most “activated”).

We use the BLOSUM90 to score every possible AA substitution. A high BLOSUM score implies that the substitution is more prevalent, which, we assume, translates into less impact on protein function. Substitutions of similar BLOSUM scores are grouped so that each group covers at least 20% of positions. Insertions and deletions, while not explicitly scored by BLOSUM, are given two separate BLOSUM group labels from all substitutions as their functional effects are assumed to be vastly different from substitutions.

Following the same reasoning that synonymous variants are assigned the same virtual “position”, they are also given the same substitution label, whose functional effect we center to 0. Also, to avoid identification problems, we force all non-synonymous AA substitution effects *ν* to sum up to 0, weighted by the count of each AA substitution group. As opposed to forcing the unweighted sum of all elements of *ν* to be 0, this weighting scheme is invariant to how AA substitutions are grouped.

This zero-sum constraint renders a simple i.i.d. prior on *ν* impossible, however. Recall that AA substitution groups are labeled 1,2, …, A where the last label is the “synonymous mutation group”. Let the mean-normalized variant count of non-synonymous mutation be *w* ∈ R^*A* —1^, then the Gaussian prior on *v* is set as follows:

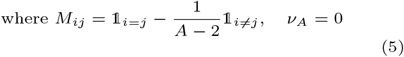

*M* is an exchangeable but degenerate correlation matrix with rank *A —* 2. *w* is mean-normalized so that if all non-synonymous mutation groups have the same number of variants, diag(w) is simply the identity matrix.

Activation scores are *ρ*_*p(υ)*_ assumed to range in [0, 1]. Intuitively, an activation score of 1 implies full activation of AA substitution effects and 0 complete inactivation. It is also assumed that such extreme activation is unusual and that most positions are moderately activated. Therefore, activation scores are given a symmetric beta prior and each *ρ*_*p(υ)*_ being independent: *ρ*_*p(υ)*_ ∼Beta(1.5,1.5)

### Aligned count error

The mean count of a variant is calculated by averaging the aligned counts *m* across time points *t* and replicates r. Variants are clustered based on such mean counts, the mapping describing which is denoted by *g*(·). The clustering is inspired by the idea that a low mean count is more susceptible to error during sampling, but `Rosace-AA` makes a weaker assumption that variants with similar mean counts have similar error scales *ϵ* in Equation 2. The prior of each error scale is independently given by: *ϵ*_*g*_*(υ)* ∼ InvGamma(1, 1)

### Numeric Bayesian inference

The hierarchical model described above is not immediately analytically tractable. Instead, we opt to conduct numeric Bayesian inference using Stan [21]. We use the default sampler offered, the No-U-Turn sampler (NUTS), which is a variant of the Hamiltonian Monte Carlo (HMC) algorithm. HMC reduces the correlation between successive samples compared to other Monte Carlo methods such as the Metropolis-Hastings Monte Carlo (MCMC) algorithm, resulting in more efficient sampling of the parameter space and thus reduced computation time for a given level of error.

Through sampling from the posterior distribution, many quantities of interest can be obtained. In the context of this paper, the functional score *β*_*υ*_of each variant can be estimated *(maximum a posteriori)* and the user may be interested in identifying variants with functional scores the most divergent from that of the wild-type.

### Rosette-AA: Family of Simulation Frameworks

In the `Rosace` paper, we introduced a data-driven growth-based DMS data simulation framework, `Rosette`, to complement real data in benchmarking the performance of `Rosace-AA` and other growth-based DMS data analysis tools. Following the pattern, we have developed a family of simulation frameworks with different sets of underlying assumptions, colloquially named Rosette-AA. to test the robustness of our models.

While `Rosette` and `Rosette-AA` have varied assumptions on how data is generated, both accept parameters that are inferred from real data. The simulation generates synthetic scores in different manners from the model, thus it serves as a robust test for the performance of models.

To avoid an unfair comparison, we designed several data simulation frameworks, based on several sets of assumptions that are increasingly complex. All frameworks begin with applying DMS analysis tools (e.g. `Rosace`, `Rosace-AA`, or OLS) to experimental data to find the functional score 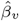 of each variant *υ*. The latent score κ_υ_ of a variant is obtained by applying a scenario-specific transformation on all variants’ functional scores. Variants are then ranked on their latent score (ties being broken randomly), labeled as negative, neutral, or positive according to their quantile, and assigned a score based on their functional identity. Here, positive and negative refer to the numeric sign - whether a negative variant is gain-of-function or loss-of-function depends on the experiment design.

The scenario-specific latent score *κ*_*υ*_ reflects the underlying assumptions of a scenario. The scenarios, ordered by ascending complexity of their assumptions, are as follows:

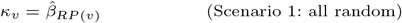

where *RP(υ)* is a random permutation of variant indices. In this simplest scenario, the scenario-specific score is the randomly shuffled functional score, so that functional identities (negative, neutral, or positive) are effectively assigned randomly.

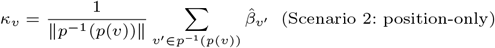

Scenario 2 assumes the effect of a variant only depends on the positional effect. Therefore, its latent score is the average of functional scores in the same position.

The last three scenarios have similar structures:

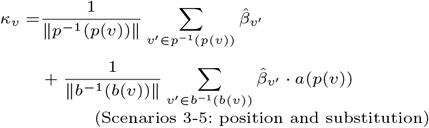

(Scenarios 3-5: position and substitution) Scenarios 3 to 5 add to the latent score the average functional scores over variants with the same amino acid substitution, with a position-level regularization term *a*(*p*(υ)) ∈ [0, 1] (i.e. “activation”). All three scenarios assume that amino acid substitution plays a role in determining the outcome.

Scenario 3 assumes *a(w) =* 1 so that there is no regularization on the mean substitution effect. It assumes that mutation effects are invariant across positions. In other words, mutation effects are assumed to be global.

Scenario 4 assumes *a(w) =* Bernoulli(0.5), allowing each position to be differently affected by mean substitution effects while assuming no further structure on the activation.

Scenario 5 assumes a deterministic activation on each position with 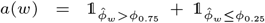. It assumes positions with very high or very low position-mean functional scores are activated differently from those with more average position-mean functional scores. Note that this formulation is statistically indistinguishable from 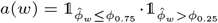.

Using the posterior distribution of each variant’s estimated functional score 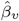, we use Bayesian hypothesis testing with its local false sign rate (LFSR) [22] to assign the estimated functional identity. In case the proportion of positive (negative) variants is too low, all positive (negative) variants will be merged into the neutral group. We have tested proteins in which all three groups can be identified, only neutral and positive groups are identified (e.g. OCT1 drug cytotoxicity screen[12]), only neutral and negative groups are identified (e.g. MET kinase domain DMS with IL3[18]), and all variants are classified as neutral (e.g. MET kinase domain DMS without IL3). We assume the variants’ functional scores of each functional identity group are Gaussian, whose parameters can be estimated using estimated functional scores of variants from that group. Variants are then ranked on their scenario-specific latent scores *κ*_*υ*._ Variants with high latent scores 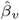 are assigned a simulated functional score 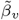 drawn from the positive distribution (if the positive functional identity group exists). The opposite applies to variants with low latent scores (if the negative functional identity group exists). The remaining variants’ simulated scores are drawn from the neutral distribution. Note that a variant with a high estimated functional score 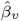 does not necessarily draw its simulated functional score 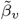from the positive distribution, as its latent score *κ*_*υ*_ can be low.

Once the simulated functional scores 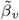 are generated for each variant, we follow the same process in Rosette to generate the simulated raw count 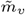.

### Secondary structure prediction model

We treat secondary structure prediction as a classification problem. To simplify, we only aim to predict *α*-helices and *β*-sheets. Each position is labeled as such, with an additional catch-all category for all other structures.

Each position has a variety of features, including raw DMS functional scores or derived statistics. We use one of the following set of features for prediction.

- `Rosace-AA`’s position-level statistics: the posterior mean of position-level functional effect *ϕ*_*p*_, error scale *σ*_*P*_, and the activation score *ρ*_*p*_.
- A vector of variant-level functional scores, indexed by which amino acid the position is mutated to (e.g., feature D of a position is the functional score of a variant that mutates the position to D, or the aspartic acid). 20-dimensional.
- Principal components of the unnormalized 20-dimensional functional scores.

We reverse the signs of some features in some data points, as protein function can have opposite effects on cell growth. We flip the sign of variant- and positional-level functional effects for experiments where loss-of-function leads to *β* >0. Other features, namely the error scale *σ* and activation score *ρ* remain unchanged.

Information on neighboring positions can help since secondary structures are contiguous. For example, if one knows both immediate neighbors of a position are part of an alpha helix, that position is likely part of the same helix. The intuition prompts the use of features from neighboring positions.

We use the SMOTE-ENN algorithm [23] to resample the training data so that the learner will not learn the *a priori* distribution of different secondary structures.

We use an ensemble of gradient boosting classifiers for classification, implemented with the XGBoost library. An XGBoost classifier is trained on the training set, using features from the neighboring *κ* positions on both sides *(2κ* + 1 in total) to predict the secondary structure of a position. The Average Precision of each classifier is calculated on the validation set, which is transformed to its weight *w*_*κ*_ = (max{AP*κ* — c,0})^2^. The weights are then normalized. The predicted probability of each class from each classifier is linearly combined using such weights. The ensemble is then retrained using both training and validation and applied to the testing dataset.

## Data and code availability

`Rosace-AA` is publicly available on GitHub (https://gitlinb.coin/ p intent el lab/r os ace-aa)

## Author contributions statement

J.R., W.C.M, and J.F. conceived the project. J.R. and M.W. designed the methods. J.R. analyzed the results. J.R. wrote the manuscript with inputs from M.H, C.M, W.C.M, and H.P. W.C.M and H.P. supervised the project.

## Acknowledgment

The authors thank the anonymous reviewers for their valuable suggestions.

